# Quantitative MHC class-I/-II gene expression levels in *CDK12* mutated prostate cancer reveal intratumorally T cell adaptive immune response in tumors

**DOI:** 10.1101/2022.04.16.487364

**Authors:** William Lautert-Dutra, Camila M. Melo, Luiz P. Chaves, Cheryl Crozier, Fabiano P. Saggioro, Rodolfo B. dos Reis, Jane Bayani, Sandro L. Bonatto, Jeremy A. Squire

## Abstract

**Background:** CDK12 inactivation is a predictive biomarker for immune checkpoint blockers (ICB) treatment response in advanced prostate cancer (PCa), but some CDK12-altered patients fail to respond to ICB. Downregulation of MHC expression has been described as a mechanism of intrinsic and acquired resistance to ICB in various cancers, but there is little information on whether MHC expression levels are altered in CDK12 defective PCa that fails to respond to ICB treatments.

**Methods:** Using genomics data of primary and metastatic prostate cancer from two public domain cohorts and a retrospective cohort, we investigated variation in the expression of the MHC genes and associated downstream changes in *CDK12* mutated patients. The findings of public domain data were validated using transcriptomic data from a 53-patient retrospective cohort from our Institute.

**Results:** Based on the analysis of gene expression quartiles, we divided the tumors into “High” and “Low” expression levels of MHC-I and -II. *CDK12* defective tumors with increased MHC levels showed the activation of several pathways associated with the immune system and elevated *PD*-*L1*, *IDO1*, and *TIM3* expression. These transcriptomic findings were confirmed using expression analyses of our cohort of 53 primary PCa. There was an increased composition of CD8+ T cells, B cells, γδ T cells, and M1 Macrophages in *CDK12* mutated tumors with elevated MHC levels based on digital cytometric analyses. In contrast, *CDK12* defective tumors with decreased MHC expression were often subject to loss of heterozygosity (LOH) genomic events affecting MHC-I/-II and the *HLA* gene cluster on chromosome 6. *CDK12* defective PCa expresses higher levels of classical MHC, has an active and inflamed tumor microenvironment, and increases the presence of effector T cells.

**Conclusions:** Reduced MHC expression may be caused by the acquisition of specific somatic genomic events that reduce the expression of antigen presentation genes. Combining CDK12 mutation, MHC expression levels, and LOH status may better predict outcomes for ICB-eligible PCa. In addition, these findings draw attention to the need to investigate therapeutic approaches for enhancing MHC expression in *CDK12* defective PCa to improve ICB responses.

## Background

Treatment of advanced prostate cancer (PCa) remains a therapeutic challenge. Men with distant metastases at diagnosis have the worst overall survival (OS), with only 30% of patients surviving > 5 years [1]. For recurrent disease, acquired resistance to androgen deprivation therapy (ADT) and chemotherapy remains a significant cause of death [2]. Immune-checkpoint blockade (ICB) therapies, such as PD-L1 inhibition, have shown only significant benefits in a minority of patients [3]. Thus, it is necessary to discover and characterize the genetic pathways and molecular signatures that could help predict more effective disease progression control in advanced PCa.

Tumor mutation burden affects the infiltration of immune cells in the tumor microenvironment (TME) [4]. Because of the high load of neoantigens, tumors with defective DNA damage repair (DDR) pathways are more suitable for immunotherapy [5]. The biallelic inactivation of Cyclin-dependent Kinase 12, *CDK12*, was recently described as a novel immune active class of advanced PCa with a more aggressive phenotype, a high mutation burden derived from focal tandem duplication events, and high inflammatory and immune cell infiltrate distinct from other defective molecular subtypes [6–9]. Although this new subtype has been proposed as a predictive biomarker of treatment response to immune-checkpoint blockade (ICB) in advanced PCa, many patients with *CDK12* alterations still fail to respond to ICB treatment. In a retrospective multi-center study, Antonarakis *et al.* found that just 33% of the *CDK12*-altered advanced PCa patients had a prostate-specific antigen (PSA) response and an increased progression-free survival of 5.4 months when treated with anti-PD-1 therapy [10]. In another study, Schweizer *et al.* showed that of the 19 advanced patients who received ICB, 11 (59%) showed a response based on a decline in PSA, with two patients (11%) having a 100% PSA decline. However, the molecular causes of ICB resistance in this molecular subtype of PCa are poorly understood [10][11].

Tumor cells may develop various escape mechanisms that allow them to avoid recognition and destruction by the immune system. For example, the expression of checkpoint proteins (PD-1/PD-L1, CTLA-4, LAG3) by tumors can modulate the activity of immune infiltrate cells [11]. They may also develop intrinsic cancer-cell signaling that can suppress infiltrating immune cells (WNT/β-catenin) and increase pro-tumorigenic immune cell infiltration (Tregs, M2 Macrophages) in the TME [12,13]. Another mechanism exploited by tumors to avoid the recognition of cytotoxic T cells (CD8+) and antigen-presenting cells (APCs) is through loss of major histocompatibility complex class-I or class-II (MHC-I and MHC-II) [14].

Changes in MHC expression have been linked to tumor progression, poor prognosis, and reduced response to ICB in different malignancies [15,16]. As classified by Garrido et al., the alteration in the MHC expression can be divided into two major mechanistic groups: tumors with “Soft” alterations are capable of recovering or upregulating MHC antigens after cytokine exposure (e.g., characterized as having regulatory abnormalities); those with “Hard” alterations cannot recover MHC expression (e.g., characterized as having structural defects) [17]. During tumor evolution, the infiltration of cytotoxic lymphocytes eliminates highly immunogenic tumor clones, causing a selection of surviving cell populations that have acquired MHC alterations through either “Soft” or “Hard” mechanistic alterations [18]. Low expression of MHC-I and -II has been associated with poor prognosis and resistance to anti-CTLA-4 and anti-PD-1, respectively [19,20]. However, in PCa, there is presently limited information on the role of MHC-I and MHC-II expression and ICB response and if the low expression of antigen presentation mechanism could better predict the lack of response to ICB in *CDK12*-altered patients.

In PCa, *CDK12* inactivation is known to increase the immunogenicity of tumor cells, but the relationship between *CDK12* loss and MHC expression has not been investigated. Changes in MHC-I and -II expression are involved in tumor immune evasion in various types of cancer. There is also evidence that higher expression of MHC genes can identify tumors likely to respond to ICB [15,16]. However, the molecular and genomic mechanisms responsible for reducing MHC expression are poorly understood. Hence, we hypothesize that variation in the expression of the MHC genes could explain the lack of response to ICB in *CDK12* defective tumors. Our transcriptomic analyses suggest that *CDK12* defective PCa expressing higher MHC levels have an active and inflamed tumor microenvironment with elevated immunomodulatory pathway expression and increased presence of effector T cells. These findings were validated using transcriptomic data from a 53-patient cohort from our Institute. Analysis of public domain data showed that tumors with decreased MHC expression also showed loss-of-heterozygosity (LOH) of MHC-I/-II and the *HLA* gene cluster on chromosome 6 associated with MHC expression and antigen presentation. Collectively, these data suggest that implementing a combined measure between *CDK12* mutation and the presence of MHC expression may better predict outcomes for prostate cancer tumors classified as eligible for ICB treatment based on *CDK12* subtype and normal antigen presentation function.

## Methods

The CbioPortal for Cancer Genomics [21] was used to search primary (pPCa) and metastatic castration-resistant (mCRPC) prostate tumors with *CDK12* mutated sample cohorts with matched clinical information. The classification of samples having *CDK12* loss-of-function followed previous reports [7–9] based only on the presence of somatic mutations (non-synonymous mutations, deep deletions, and shallow deletions) in both *CDK12* alleles (*CDK12* defective) [6–8]. Only cohorts containing *CDK12* defective samples WES and RNAseq data were selected in the Genotypes and Phenotypes (dbGaP) database and applied for access under project ID 29255 (Additional file 1: Figure S1, additional table 1).

**Table 1.** Enrichment results. Enriched GO (BP) pathways of the upregulated genes from TCGA CDK12-Mut MHC-I and MHC-II High, mCRPC group MHC High, and for our validation cohort (RP) MHC-I High group. Enrichment analyses were performed using clusterProfiler and p-adjusted value = 0.05 as the cutoff.

|  | ID | Description | p.adjust |
| --- | --- | --- | --- |
| TCGA-PRAD_MHC-I | GO:0051249 | regulation of lymphocyte activation | 1,48E-19 |
|  | GO:0002460 | adaptive immune response based on somatic recombination of immune receptors built from immunoglobulin superfamily domains | 1,48E-19 |
|  | GO:0050867 | positive regulation of cell activation | 2,78E-20 |
|  | GO:0002449 | lymphocyte mediated immunity | 1,16E-18 |
|  | GO:0006958 | complement activation, classical pathway | 2,77E-14 |
|  | GO:0006959 | humoral immune response | 1,08E-09 |
|  | GO:0050853 | B cell receptor signaling pathway | 2,20E-08 |
|  | GO:0002440 | production of molecular mediator of immune response | 7,15E-08 |
|  | GO:0042742 | defense response to bacterium | 1,39E-07 |
|  | GO:0006910 | phagocytosis, recognition | 5,66E-08 |
| TCGA-PRAD_MHC-II | GO:0051249 | regulation of lymphocyte activation | 2,00E-25 |
|  | GO:0042110 | T cell activation | 5,84E-25 |
|  | GO:0050867 | positive regulation of cell activation | 9,60E-23 |
|  | GO:0007159 | leukocyte cell-cell adhesion | 2,50E-20 |
|  | GO:0002460 | adaptive immune response based on somatic recombination of immune receptors built from immunoglobulin superfamily domains | 2,80E-20 |
|  | GO:0002443 | leukocyte mediated immunity | 3,53E-19 |
|  | GO:1903037 | regulation of leukocyte cell-cell adhesion | 8,17E-19 |
|  | GO:0070661 | leukocyte proliferation | 1,75E-15 |
|  | GO:0046651 | lymphocyte proliferation | 5,77E-14 |
|  | GO:0070663 | regulation of leukocyte proliferation | 1,64E-12 |
| mCRPC_MHC | GO:0051251 | positive regulation of lymphocyte activation | 4,87E-15 |
|  | GO:0002449 | lymphocyte mediated immunity | 6,43E-15 |
|  | GO:0002253 | activation of immune response | 6,67E-15 |
|  | GO:0042110 | T cell activation | 3,27E-11 |
|  | GO:0002460 | adaptive immune response based on somatic recombination of immune receptors built from immunoglobulin superfamily domains | 1,76E-09 |
|  | GO:0006959 | humoral immune response | 1,21E-06 |
|  | GO:0019724 | B cell mediated immunity | 2,55E-06 |
|  | GO:0050863 | regulation of T cell activation | 9,19E-07 |
|  | GO:0007159 | leukocyte cell-cell adhesion | 3,27E-04 |
|  | GO:1903037 | regulation of leukocyte cell-cell adhesion | 1,75E-03 |
| RP_MHC | GO:0019221 | cytokine-mediated signaling pathway | 1,32E+11 |
|  | GO:0022409 | positive regulation of cell-cell adhesion | 1,32E+11 |
|  | GO:0002474 | antigen processing and presentation of peptide antigen via MHC class I | 1,21E+12 |
|  | GO:0072182 | regulation of nephron tubule epithelial cell differentiation | 1,21E+12 |
|  | GO:0060333 | interferon-gamma-mediated signaling pathway | 0.001 |
|  | GO:0002479 | antigen processing and presentation of exogenous peptide antigen via MHC class I, TAP-dependent | 0.001 |
|  | GO:0042127 | regulation of cell population proliferation | 0.001 |
|  | GO:0042590 | antigen processing and presentation of exogenous peptide antigen via MHC class I | 0.001 |
|  | GO:0010719 | negative regulation of epithelial to mesenchymal transition | 0.002 |
|  | GO:0048640 | negative regulation of developmental growth | 0.002 |

We used a retrospective cohort of clinical intermediated primary PCa derived from radical prostatectomies (*n*=53) to validate our transcriptomic findings. All 53 samples included in the cohort were primary PCa collected by radical prostatectomy following National Comprehensive Cancer Network (NCCN) clinical practice guidelines [22] in the Department of Surgery and Anatomy, Urology Division at Ribeirao Preto Medical School, Brazil, between 2007 and 2015 (Supplementary Table S1). The patients were classified according to the presence of biochemical recurrence (BCR), defined as PSA>0.2 ng/ml within six months after radical prostatectomy. Patient outcome data were collected to the last follow-up date (additional table 1). This retrospective study was approved by the Ethics Committee in Research of Hospital of Ribeirão Preto, São Paulo, Brazil (HCRP) number CAAE 60032122.8.0000.5440 and the Ethics Board of the University of Toronto (Protocol: 00043323).

### RNA/DNA isolation

The RNA was isolated from tissues with tumor-rich areas previously marked by a pathologist (FPS). Sections were processed at the Ontario Institute for Cancer Research, Toronto, Canada (OICR) using a dual DNA and RNA extraction as previously described [23,24]. Briefly, hematoxylin and eosin slides were prepared for all the Formalin-Fixed Paraffin-Embedded (FFPE) tissues. Histologic analysis of all the slides was performed in the pathology department, and all tumor areas were carefully marked by an experienced pathologist (FPS). The percentage of tumor cells (range 70-95% tumor-rich) within each marked tumor-rich area was estimated and recorded (Table S2). Adjacent slides for each tumor were prepared, and the same areas of interest were microdissected for RNA extraction.

**Table 2.**
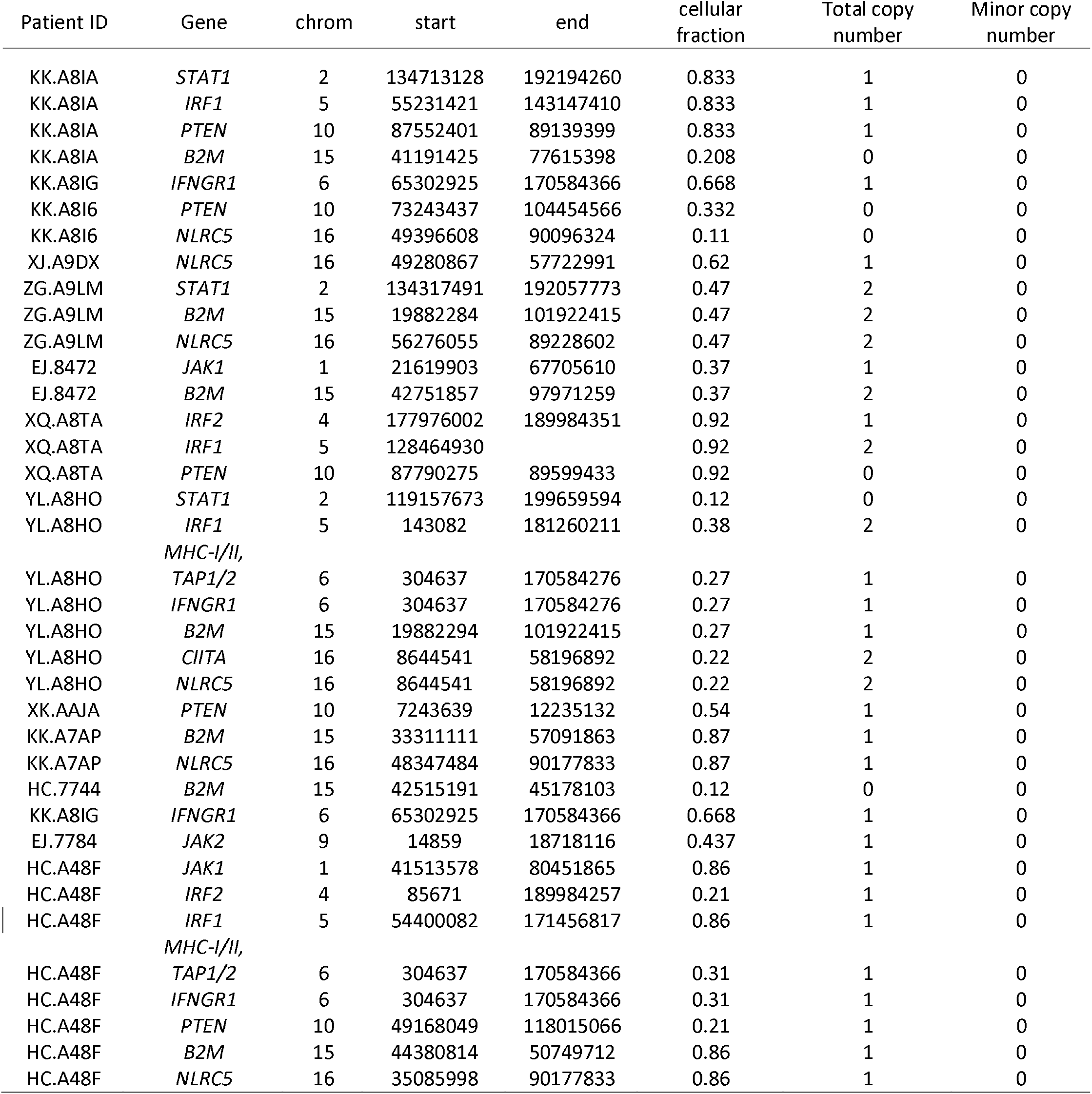
CDK12 defective in primary MHC low expressed tumors. Integral results of FACETS analysis of whole-exome sequencing data from CDK12 defective primary tumors expressing low levels of MHC-I/II. The table shows the integer copy number for the allelic ratio of the heterozygous SNPs in each patient’s tumor/normal pair. The first column is the patient ID, and the second column is the MHC region loci and other genes commonly subject to LOH in PCa. The last two columns represent the total copy number and the minor copy number, respectively. Tumors with a 2:1 ratio are considered normal for each position, while 2:0 and 1:0 represent CN-LOH and LOH events. The cellular fraction represents the estimated number of cells harboring the genotype. Primary PCa revealed subclonal CN-LOH and LOH at JAK1, STAT1, IRF2, IRF1, MHC-I/II, TAP1, TAP2, IFNR1, JAK2, B2M, CIITA, NLRC5, and IFNR1. Two patients showed biallelic loss at the B2M locus (Total copy number/minor copy number ratio=0:0). *MHC-I = HLA-A, HLA-B, and HLA-C; MHC-II = HLA-DRA, HLA-DRB5, HLA-DRB6, HLA-DPA1, HLA-DPB1, HLA-DQA1, HLA-DQB1, and HLA-DQB2.

### Transcriptomic analysis

For the pPCa (TCGA-Prostate Adenocarcinoma, *n* = 48), we used the package recount2 to download summarized experiments objects containing the transcription-level RNA-Seq abundance matrix [25,26]. For the mCRPC (SU2C, *n* = 10), we downloaded SRA Paired-End (PE) reads using the SRAToolkit. The quality of the raw reads was then measured using the FASTQC program. We quantified the transcripts using the software Salmon v1.6.0 [27]. The transcriptome index was built using the reference GRCh38 version of the human genome and transcriptome, downloaded from ENSEMBL and GENCODE, following the manual instruction. We then used the Tximport v1.22.0 package to import the transcription-level abundance and estimate raw counts derived from the quantification step. The count data normalization, expression levels, and differential gene expression (DEG) analysis for pPCa and mCRPC cohorts were executed using the DESeq2 v1.34.0 [28,29]. We used the clusterProfiler v4.2.1 to implement equally over-representation (ORA) enrichment analysis and gene set enrichment analysis (GSEA) of the DEGs and the whole transcriptome profile, respectively, using the Gene Ontology (GO) and Molecular Signatures Database (MSigDB) [30–35] (Figure S1).

The PanImmune Panel (NanoString Technologies Inc., Seattle, WA, USA) was used to profile the RNA derived from the FFPE samples from our 53-patient tumor cohort. Raw expression data from the PanImmune Panel was loaded in nSolver software v4.0 (NanoString Technologies) to perform the quality control (QC analysis) and to build the transcript matrix for downstream analysis.

### MHC expression

We performed a hierarchical clustering analysis using representative MHC genes. We observed two groups of *CDK12* defective tumors based on MHC expression, as shown in Additional file 1, Figure S1. We dichotomized expression levels below or above the first quartile for each gene; we then used this classification to generate the final logical values regarding the patient’s MHC status (‘High’ or ‘Low’ expressed) (Additional file 1: Figure S2)[36,37]. For the quartile quantification, we used the normalized expression values of the classical genes that composed each MHC class (e.g., MHC-I: *HLA-A*, *HLA-B*, *HLA-C*; MHC-II: *HLA-DPA1*, *HLA-DPB1*, *HLA-DQA1*, *HLA-DQB1*, *HLA-DQB2*, *HLA-DRA*, *HLA-DRB5*, *HLA-DRB6*) [37]. Samples were classified as having MHC ‘Low’ expression when at least one gene composing each class was expressed at a low level. In mCRPC *CDK12*-defective, all tumors classified as MHC-High presented similar high expression of both MHC-I and -II; thus, we called these tumors just MHC High (henceforth, mCRPC *CDK12* defective MHC High). The summary values used to classify the tumors are shown in table S1. Our retrospective cohort of low-intermediate risk primary PCa derived from radical prostatectomies was used to validate our transcriptomic findings that were derived from public domain cohorts.

### Genomic profile

We downloaded SRA Paired-End (PE) reads for both pPCa (n=48) and mCRPC (n=10) and processed it as previously described. The GRChg38 reference was sorted using the SeqKit [38]. The final fastq files were then aligned to hg38 using bwa with a penalty for up to 3 mismatches per read [39,40]. Sam files were converted to bam files and processed using samtools v1.16.1. To determine whether any MHC expression differences were related to genomic alterations, we used the FACETS (Fraction and Allele-Specific Copy Number Estimated from Tumor Sequencing) algorithm [41]. This approach uses matched normal-tumor WES and provides mutant allele-specific copy-number homozygous/heterozygous deletions, copy-number neutral loss-of-heterozygosity (LOH), allele-specific gain/amp in genomic loci associated with genes involved in antigen presentation (Additional table 2). Reference and variant allele read counts were extracted from the bam file for common, polymorphic SNPs downloaded from dbSNP (GRCh38p7) using FACETS *snp*_*pileup* function with a minimum threshold for mapping quality, the minimum threshold for the base quality, and minimum read depth of 15, 20, 20, respectively. The pre-processing followed the suggested recommendations from the manual, and genomic intervals of 150-250bp were used to avoid hyper-segmentation in high polymorphic neighborhood regions. Mutant allele-specific copy-number changes were declared when the points changes were greater than a pre-determined critical value (cval) of 100 compared to constant copy-number regions [42,43].

### Digital cytometry

To investigate and quantify the immune cell composition in the TME of tumors having *CDK12* defects, we used the bulk tissue gene expression profiles (GEP) from RNA-seq data from both pPCa and mCRPC with the digital cytometry CIBERSORTx [44–46]. CIBERSORTx uses bulk tissue GEP, compares the data with prior knowledge of expression profiles from purified leukocytes, and estimates a tumor’s relative immune abundance composition. We used the ‘signature matrix’ containing a validated leukocyte GEP of 22 human hematopoietic cell phenotypes, leukocyte gene signature matrix (LM22), to estimate the immune cell composition from the TME.

### Computational and statistical analysis

A GNU/Linux environment was used to perform the quality control and quantify the raw reads to the human transcriptome. Subsequently, downstream analysis was performed in RStudio software (R Foundation for Statistical Computing, R v4.1.2). Pearson Correlation was used to analyze the normalized expression levels (coef. level = 0.95). A gene was considered differentially expressed when log2 foldchange > 1 was expressed from the reference group with a *p*-adjusted value < 0.05. For the enrichment analysis, we used a cutoff value of 0.05.

## Results

### Transcriptomics

We hypothesize that variation in the expression of the MHC genes among *CDK12* defective tumors could provide predictive information on poor ICB responses in this molecular subtype of PCa. To investigate MHC expression levels in PCa further, we compared the transcriptomics of prostate tumors, including *CDK12* defective (*CDK12* defective) prostate tumors, classified as “MHC High” (top 75% quartile) to the transcriptomics of tumors classified as having a low expression of MHC genes (bottom 25% quartile).

### CDK12 defective MHC-I/-II High-expression tumors showed distinct transcriptome profiles with upregulation of IFN-γ-responsive genes and an inflamed TME

We first compared the transcriptome of *CDK12* defective tumors classified as MHC-High and MHC-Low (Figure 1a-c, Additional file 1: Figure S2). The 48 primary PCa *CDK12* defective MHC-I High showed 1319 upregulated and 2153 downregulated genes (Additional table 3, 4) and showed four apparent clusters (Figure 1a). The upper cluster (I) demonstrates *IGHV* and *IGLV* over-expressed genes linked to *CDK12* MHC I high. The central clusters (II, III) have four *HLA* genes overexpressed in the MHC I high group, which supports our classification based on *CDK12* defective MHC expression (e.g., *HLA-A*, *HLA-B*, *HLA-C*). Also, this cluster showed upregulation of many genes related to antigen presentation and CD8+ T cell activity (Figure 1a). The lower cluster (IV) possesses 14 downregulated genes when MHC I is highly expressed. Likewise, *CDK12* defective MHC-II High exhibited 1129 upregulated and 580 downregulated genes (Additional tables 3, 5). Our comparison showed two clusters (Figure 1b). The upper cluster showed the upregulation of genes related to cytotoxicity, immune cell migration, and immune suppression (Figure 1b).

**Figure 1.**
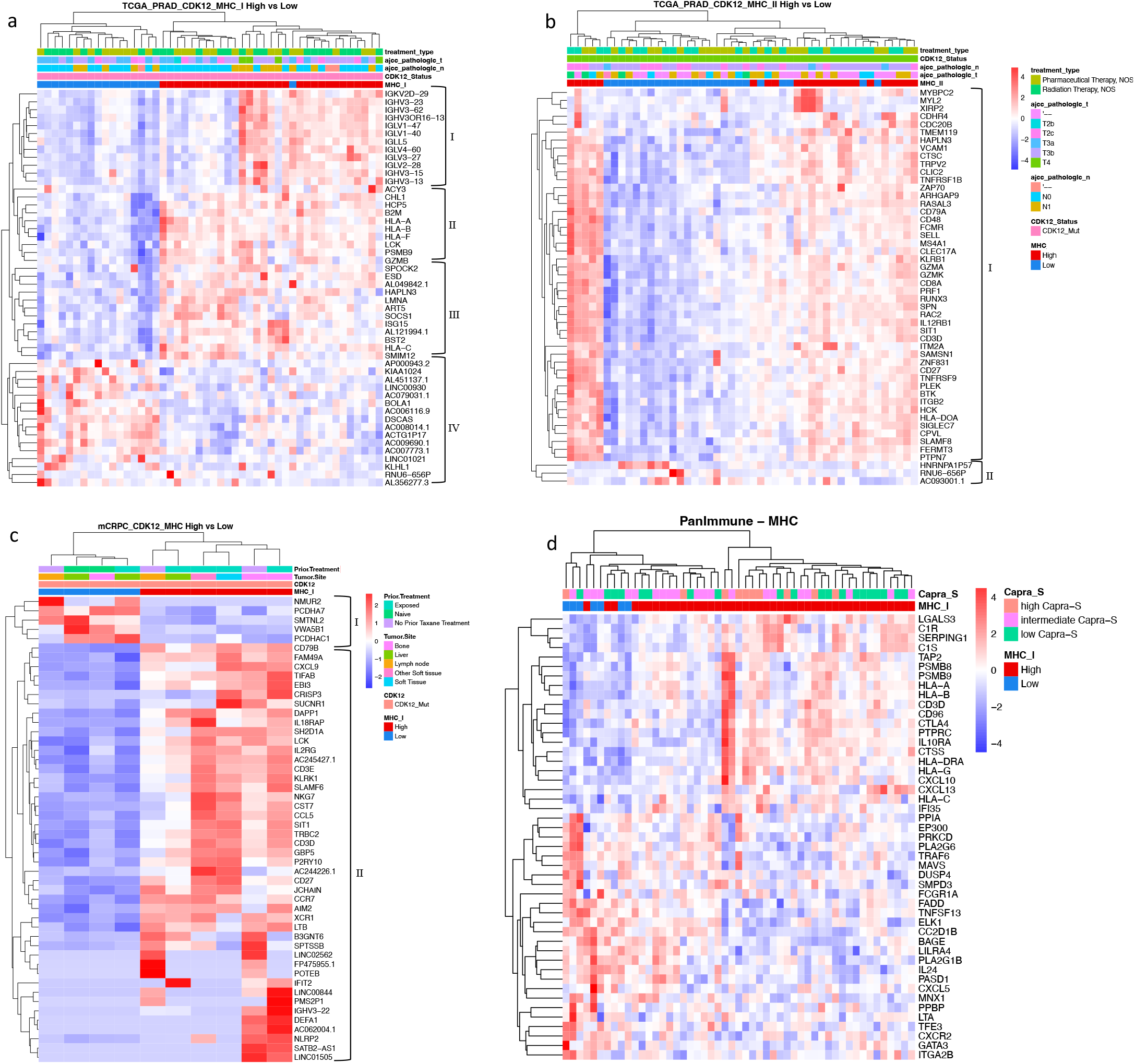
MHC-I/-II high expression in CDK12 defective prostate cancer showed distinct transcriptome activity. **(a)** Transcriptome heatmap exhibiting clustering of top 50 DEGs in pPCa CDK12-Mut MHC-I High group (*n* = 30). The upper cluster (I) has IGHV and IGLV over-expressed genes linked to *CDK12* MHC I high. The central clusters (II and III) have four HLA genes overexpressed in the MHC I high group, supporting our CDK12 mutated MHC expression classification. The lower cluster (IV) has 14 downregulated genes when MHC I is highly expressed. **(b)** Transcriptome heatmap exhibiting clustering of top 50 DEGs in the pPCa CDK12-Mut MHC-II High group (*n* = 23). The upper cluster (I) has upregulation of genes related to cytotoxicity, immune cell migration, and immune suppression. We could not observe clinical features associated with MHC I or II clusters. **(c)** Transcriptome heatmap exhibiting clustering of top 50 DEGs in the mCRPC CDK12-Mut MHC High group (*n* = 6). The combined MHC I and II showed two distinct clusters. The lower cluster showed the upregulation of genes linked to innate and adaptive immune response and CD4+ and CD8+ T cell activity. **(d)** Transcriptome heatmap exhibiting clustering of the DEGs from our validation cohort (RP cohort). The DEGs are relative to the CDK12-Mut ‘Low’ group. Clinical information is displayed on top of the heatmap for each patient. The color scale in the heatmap represents the Z-score of the normalized read counts for each gene, where the red scale indicates upregulated and blue low-expressed genes.

The analysis of ten mCRPC tumors identified 230 upregulated and 46 downregulated genes differentially expressed in *CDK12* defective MHC-high (Additional tables 3, 6) and showed two distinct clusters (Figure 1c). The lower cluster showed the upregulation of genes linked to innate and adaptive immune response CD4+ and CD8+ T cell activity (Figure 1c). In conclusion, for both primary PCa and metastatic CRPC, *CDK12* defective PCa that express higher MHC levels was associated with transcriptomic changes that indicate a general pattern of activation of IFN-γ-responsive and cytotoxic activity genes.

In our institutional validation cohort, we identified 25 upregulated and 14 downregulated genes that were differentially expressed in the MHC-high group and showed two distinct clusters (Figure 1d). The patients classified as MHC I high group have *HLA* genes overexpressed, consistent with our classification based on RNA-seq using both public domain cohorts. Thus, for both primary PCa and metastatic CRPC, *CDK12* defective PCa that express higher MHC levels was associated with transcriptomic changes that indicate a general pattern of activation of IFN-γ-responsive and cytotoxic activity genes.

To better understand the transcriptional alterations in *CDK12* defective tumors classified as MHC-I/-II high-expressed, we used ORA and GSEA analysis to identify potential functional associations of expression changes associated with differentially expressed genes. In primary PCa, tumors with *CDK12* defective MHC-I and MHC-II high-expressed showed significative enrichment of pathways related to activation of immune cells and antigen presentation by MHC-I and MHC-II, as expected (Table 1, Additional table 4, 5). Furthermore, GSEA results showed activation of many pathways in *CDK12* defective MHC-I high-expressed (e.g., Interferon Gamma Response, Interferon Alpha Response, TNFA Signaling via NFKB, and IL2/STAT5 Signaling (Figure 2a, Additional table 4). In addition, the *CDK12* defective MHC-II High group had similar pathway activation but with the addition of Allograft Rejection (Figure 2b, Additional table 5).

**Figure 2.**
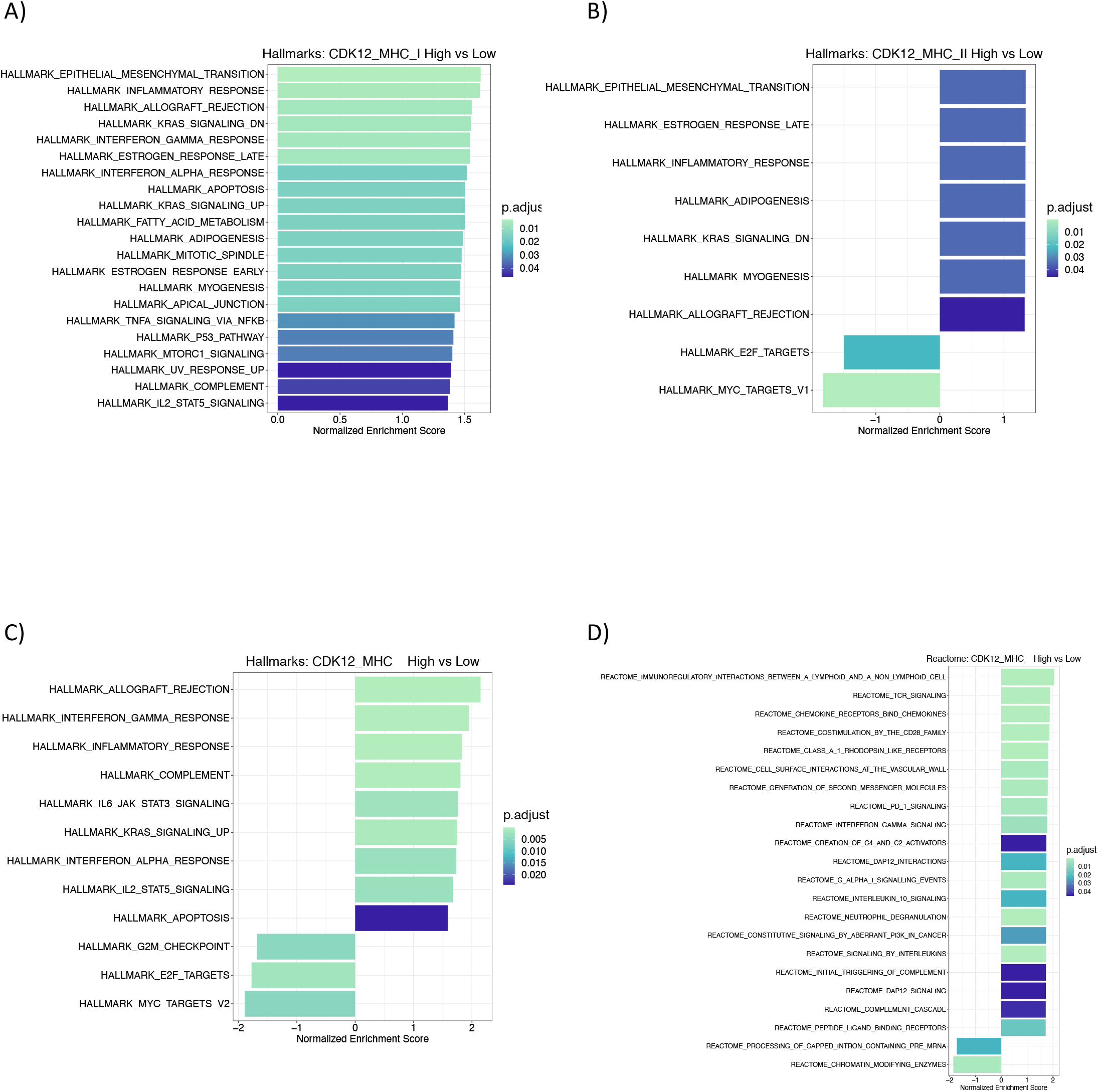
MHC high expressed *CDK12* defective prostate tumors are associated with activation of immune-related pathways. **(a)** Gene Set Enrichment Analysis of Hallmark’s pathways in pPCa *CDK12-Mut* MHC-I High (*n* = 30)**. (b)** Gene Set Enrichment Analysis of Hallmark pathways in pPCa *CDK12-Mut* MHC-II High (*n* = 23). **(c-d)** Gene Set Enrichment Analysis of Hallmarks and Reactome pathways in mCRPC *CDK12-Mut* MHC High (*n* = 10). A normalized enrichment score indicates each factor’s positive or negative association with the condition of interest, which means activation or suppression of the pathway. Enrichment analyses were performed using *clusterProfiler* and *p*-adjusted value = 0.05 as the cutoff.

The *CDK12* defective metastatic group with MHC high-expressed showed enrichment of cytotoxicity and adaptive cytotoxicity immune response (Table 1, Additional table 6). In addition, GSEA results showed activation of various hallmark and Reactome pathways (e.g., Allograft Rejection, Interferon Gamma Response, Interferon Alpha Response, IL2/STAT5 Signaling, TCR Signaling, Interleukin 10 Signaling, and the suppressive PD1 Signaling pathway) (Figure 2c-d, Additional table 6).

Next, to investigate and understand the involvement of the DEGs in pathways driving tumor progression, we further used ORA analysis on the DEGs from our validation cohort. As expected, tumors with MHC-I high-expressed showed activation of several immune-related pathways (Table 1, Additional file 1: Figure S4).

Our transcriptomic analysis suggests that *CDK12* defective PCa expressing high MHC genes display the activation of various pathways associated with an active and inflamed tumor microenvironment. Also, these results demonstrate the feasibility of using MHC expression levels to uncover intrinsic inflammatory activity in PCa.

### CDK12 defective MHC-I/-II High groups exhibited tumor microenvironment with high expression of chemokines and immunomodulatory genes

We investigated the difference between chemokines and immunomodulatory gene expression to elucidate the underlying immunologic signature in the TME of CDK12 defective tumors expressing high MHC genes. In primary PCa, the *CDK12* defective MHC-I High group exhibited upregulation of many chemokines linked to APCs and effector T cell migration (Addition file 1, Figure S4) and the immunomodulatory genes *HAVCR2* (*TIM3*) and *IDO1*. (Figure 3a-b). Also, the MHC-II high group presented increased expression of chemokines related to Natural killer, T effector cell, and B cell infiltration (Addition file 1, Figure S2, Figure S5), and the immunomodulatory genes *CD274* (*PD-L1*), *HAVCR2* (*TIM3*), and *IDO1* (Figure 3c-e). Based on Pearson Correlation analysis, we observed a significant positive correlation between MHC-I complex genes and *IDO1* but with no other investigated gene (Addition file 1, Figure S7a). Interestingly, among the MHC-II genes, correlation analysis showed a significant positive association between the expression of the MHC-II complex and the immunomodulatory genes *CD274* (*PD-L1*), *IDO1*, and *HAVCR2* (*TIM3*) (Figure S7b).

**Figure 3.**
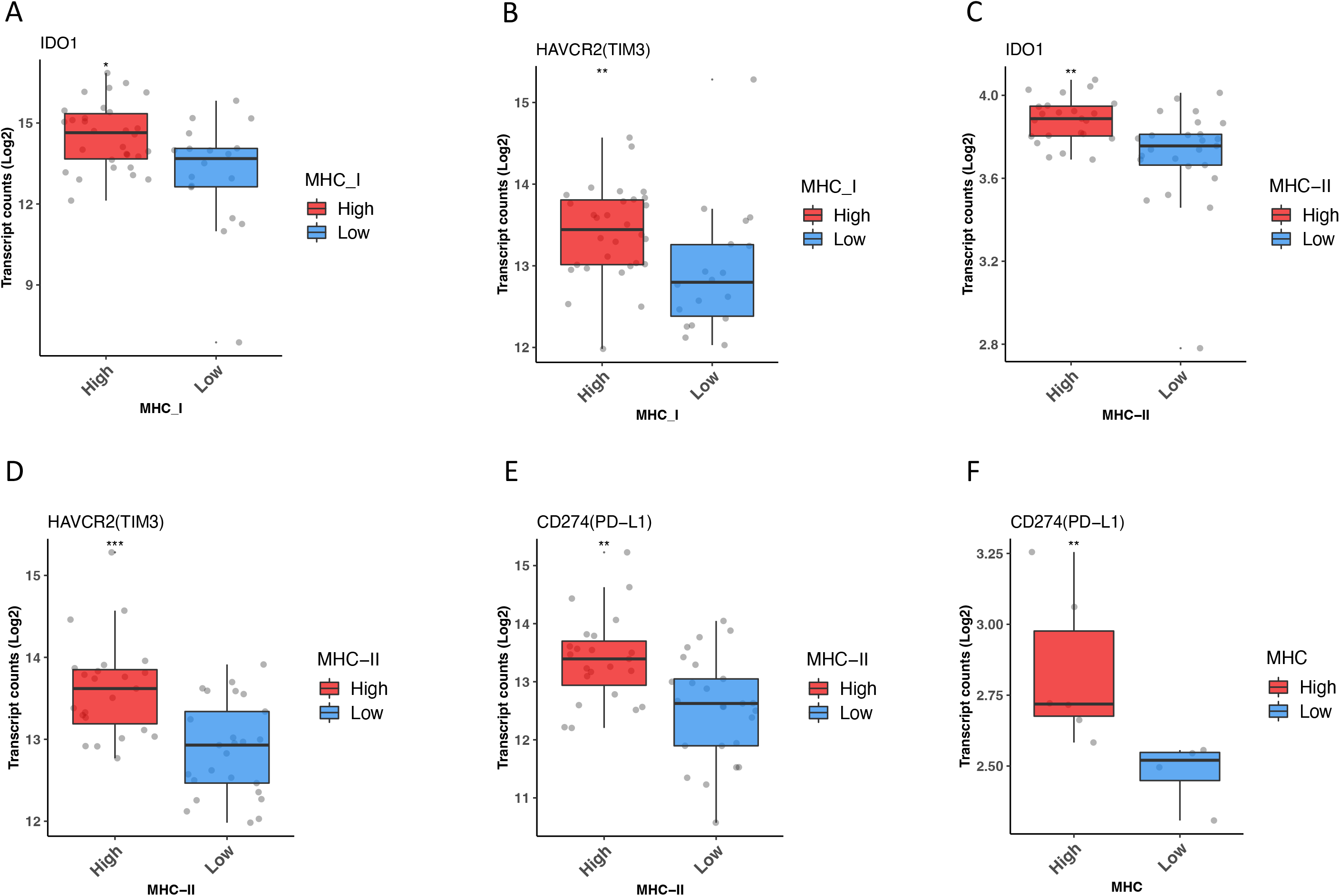
MHC high expressed CDK12 defective prostate cancer showed enhanced expression of immunomodulatory genes. RNA-sequencing expression level of immunomodulatory genes **(a)** *IDO1* (*p* < 0.018), **(b)** *HAVCR2* (*TIM3*) (*p* < 0.0022) in *CDK12-Mut* MHC-I High. *n* = 48; **(c)** *CD274* (*PD-L1*) (*p* < 0.0028), **(d)** *HAVCR2* (*TIM3*) (*p* < 0.00012), **(e)** *IDO1* (*p* < 0.001) in *CDK12-Mut* MHC-II High. *n* = 48; **(f)** *CD274* (*PD-L1*) (*p* < 0.0095)*. n* = 10; *: *p*-value < 0.05; **: *p*-value < 0.01, by Mann-Whitney test. *IDO1*, Indoleamine 2,3-Dioxygenase 1; *HAVCR3* (*TIM3*), T-Cell Immunoglobulin Mucin Receptor 3; *CD274* (*PD-L1*), Programmed Cell Death 1 Ligand 1.

The *CDK12* defective metastatic group with high expression of MHC showed high expression of chemokines (Addition file 1, Figure S6) and the immunomodulatory gene *CD274* (*PD-L1*) (Figure 3f). Furthermore, we observed a significant positive association between the immunomodulatory genes *HAVCR2* (*TIM3*) and *CTLA4*, *LAG3*, and MHC-I and -II genes (Addition file 1, Figure S7c-d). *CDK12* defective tumors’ expression of higher levels of MHC genes shows a pattern of upregulation of chemokine and immunomodulatory mechanisms.

### CDK12 defective MHC-I/-II High CDK12 defective tumors showed distinct immune cell infiltrate in the TME

Our transcriptomics analysis in the public domain cohort predicted that *CDK12* defective tumors expressing high levels of MHC-I/-II genes would display enrichment of immune cells with high effector and cytotoxic activity and the co-inhibitory expression of inhibitory pathways by tumor cells (Figures 1, 2, 3). We used *in silico* cytometry (CIBERSORTx) to test this hypothesis and to estimate the immune cell composition in the TME of *CDK12* defective tumors expressing high levels of MHC-I/-II genes.

We investigated whether tumors expressing high MHC-I and -II levels affected the variation in immune cell composition in *CDK12* defective tumors in both public domain cohorts. In primary PCa, *CDK12* defective MHC-I high-expressed showed a significant increase in the composition of Naïve B cells, CD8+ T cells, and γδ T cells. In contrast, the reduced composition of Mastocytes (Figure 4a). In comparison, the MHC-II high-expressed group exhibited an increase of γδ T cells and reduced composition of Plasma cells and M0 macrophages (Figure 4b).

**Figure 4.**
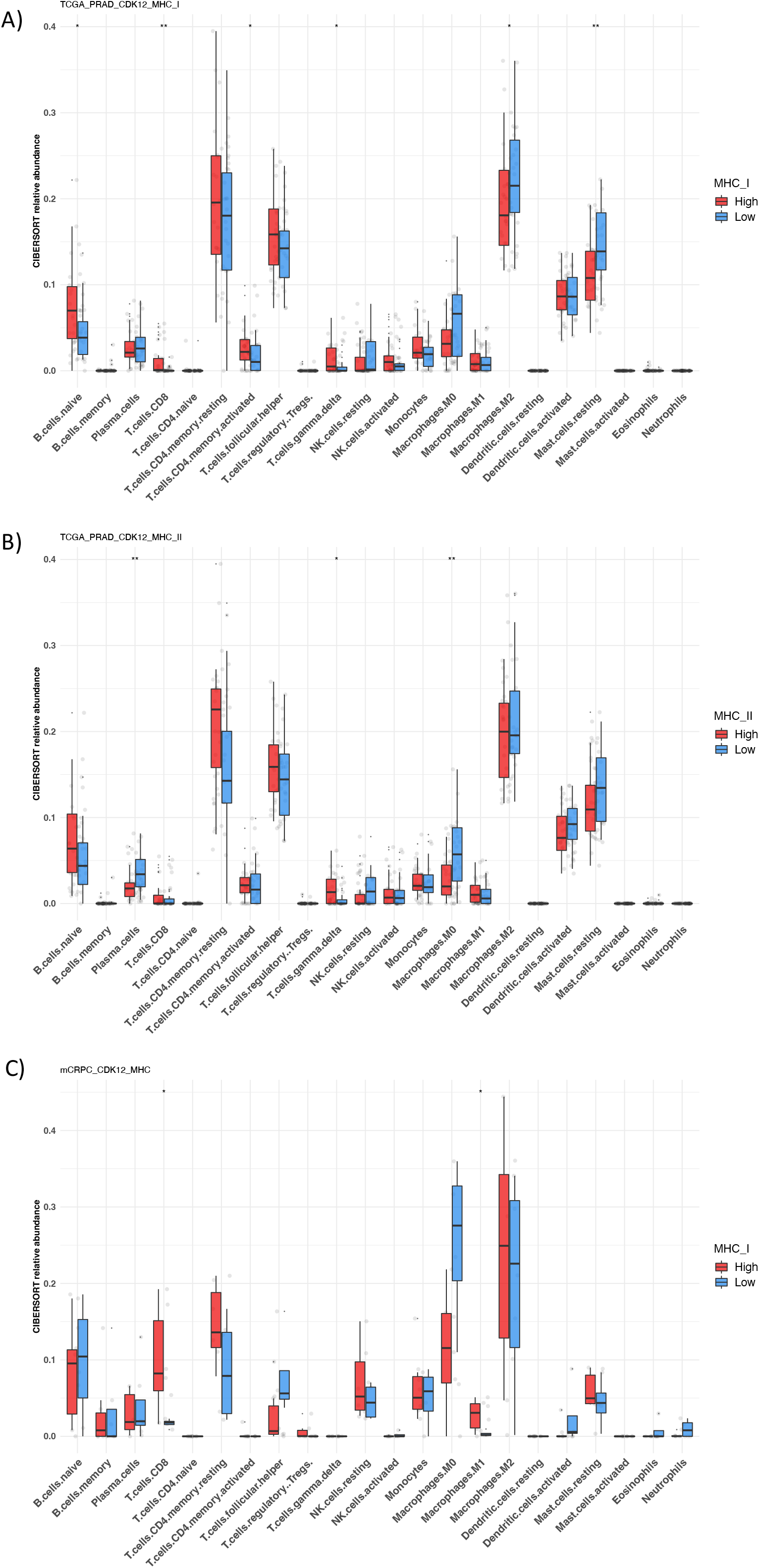
MHC high expressed *CDK12* defective is associated with enhanced T cell recruitment of prostate tumors. **(a)** CIBERSORT-derived immune cell abundance of 22 distinct cell subsets based on *CDK12* defective MHC-I High pPCa (*n* = 30)**. (b)** CIBERSORT-derived immune cell abundance of 22 distinct cell subsets based on *CDK12* defective MHC-II High pPCa (*n* = 23). **(c)** CIBERSORT-derived immune cell abundance of 22 distinct cell subsets based on *CDK12* defective MHC High mCRPC (*n* = 10). *: *p*-value < 0.05; **: *p*-value < 0.01, by Mann-Whitney test.

Additionally, the *CDK12* defective metastatic group with high expression of MHC showed a significative increased composition of CD8+ T cell and M1 Macrophages (Figure 4c). Our results suggest that *CDK12* defective PCa expressing higher levels of MHC displayed increased APCs and effector lymphocytes traffic in their TME.

### Genomic alterations of MHC-I/-II region associated with defective CDK12

During tumor evolution, the infiltration of cytotoxic lymphocytes eliminates highly immunogenic tumor clones, causing a preferential selection and survival of cell populations that have acquired MHC alterations. To determine whether the low levels of MHC-I/-II genes in *CDK12* defective tumors were derived from somatic genomic alterations affecting antigen presentation genes, we used an allele-specific copy number algorithm to estimate the copy number profile of *CDK12* defective tumors expressing low levels of MHC-I/-II genes. The algorithm incorporates quantitative analysis of RNA-seq data derived from MHC-I/-II and *HLA* genes on chromosome 6 and several other genes, such as *PTEN*, commonly subject to LOH in PCa [47]. Estimates include the determination of the relative clonality of allele-specific copy number alterations based on the fraction of tumor cells bearing LOH.

WES of primary PCa revealed both subclonal allele-specific copy-neutral losses of heterozygosity (CN-LOH) and complete loss of heterozygosity (LOH) events affecting MHC-I/-II and *HLA* genes on chromosome 6. Both major clonal and minor subclonal events were detected by comparing the copy number to the estimated cellular fraction of tumor cells harboring the copy number alteration (Table 2 and Figure S8). One example is illustrated by tumor TCGA-KK-A8IA in which only one major clonal event was detected with a cellular fraction = 0.833, and LOH events were detected on chromosomes 2, 5, and 10. While TCGA-YL-A8HO revealed four subclonal events, capturing biallelic loss, CN-LOH, and LOH in chromosomes 2, 5, 6, 15, and 16.

Tumors expressing lower levels of MHC-I/II genes may be associated with the somatic genomic LOH events affecting regional transcriptional regulators of MHC expression and the cis-acting regulatory components of the MHC (Table 2). Two patients also showed subclonal complete loss of the *B2M* locus (ID: TCGA-EJ-8471, cellular fraction = 0.37; TCGA-YL-A8HO, cellular fraction = 0.27. Table 2). Interestingly, *B2M* is an important component of the MHC-I complex, and mutations of this gene have previously been associated with ICB resistance [48]. In contrast to primary PCa, metastatic tumors harboring *CDK12* mutation expressing low levels of MHC-I/II showed allele-specific copy number gains in the MHC-I/I*I* complex and key regulators of MHC expression (Table 3 and Figure S9). Interestingly, one patient showed a clonal LOH event at the *JAK2* and *B2M* loci (ID: 5115615, cellular fraction = 0.90).

**Table 3.**
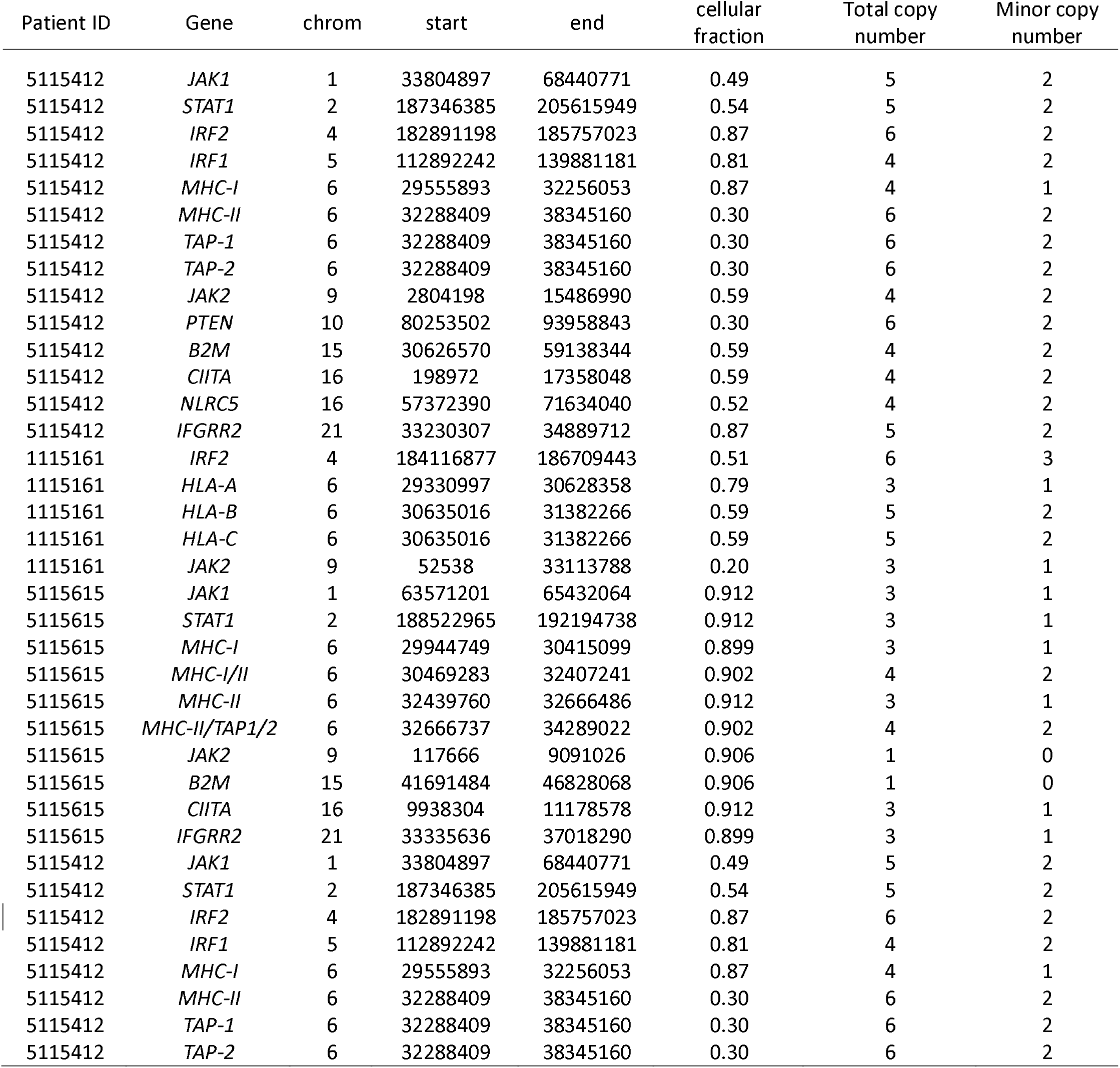
*CDK12* defective in metastatic MHC low expressed tumors. Integral results of FACETS analysis of whole-exome sequencing data from *CDK12* defective metastatic tumors expressing low levels of MHC-I/II. The same parameters shown in Table 1 were used to characterize LOH for each gene of interest. Metastatic PCa showed allele-specific copy number gains at *JAK1, STAT1, IRF2, IRF1, MHC-I/II, TAP1, TAP2, IFNR1, JAK2, B2M, CIITA, NLRC5,* and *IFNR1*. One patient showed a clonal LOH event at the *JAK2* and *B2M* loci (Total copy number/minor copy number ratio=1:0). *MHC-I = *HLA-A, HLA-B,* and *HLA-C;* MHC-II = *HLA-DRA, HLA-DRB5, HLA-DRB6, HLA-DPA1, HLA-DPB1, HLA-DQA1, HLA-DQB1,* and *HLA-DQB2*.

These data indicate that low expression of MHC in primary tumor clones appears to be associated with frequent somatic CN-LOH and LOH subclonal genomic alterations in primary tumors harboring *CDK12* mutation. While metastatic tumors revealed both LOH and high copy-number gains affecting antigen presentation genes and their regulators. Therefore, these subclonal CN-LOH and LOH events are associated with impaired MHC expression that may contribute to reducing tumor immunogenicity.

## Discussion

In PCa, *CDK12* inactivation is known to increase the immunogenicity of tumor cells [9,49–52]. MHC-I and -II expression changes are involved in tumor immune evasion in various cancer types [37,53–55]. Using *in silico* two public domain RNA-seq cohorts and transcriptional profiling of our institutional cohort, we observed two groups of *CDK12* defective tumors regarding MHC expression. Tumors expressing higher levels of MHC showed the activation of multiple pathways associated with the immune system, significantly high expression of immunomodulatory genes, and increased CD8+ T cells, B cells, γδ T cells, and M1 Macrophages composition in their TME. Also, Tumors with low MHC expression showed allele-specific copy-number alteration in genes involved in regulating MHC expression and antigen presentation.

Analysis of the clinical data showed that higher expression of MHC genes in pPCa and mCRPC, respectively, exhibited a trend towards better overall survival when compared to the MHC lower expression group (*p* > 0.05, Figure S3). Several studies have found similar MHC-I and -II gene expression relationships associated with better clinical outcomes and different resistance mechanisms [20,56,57]. Lack of MHC-I expression in melanoma was linked to higher odds of progressive disease and resistance to anti-CTLA4 but not anti-PD1 [20]. In the same study, expression of MHC-II in tumors was associated with tumors more likely to respond to anti-

PD1 treatment. MHC-II expression was linked to a positive outcome in Hodgkin Lymphoma treated with nivolumab (anti-PD1) [56]. Park et al. showed that MHC-II positivity in tumor cells was associated with better disease-free survival in patients with lymph node metastasis [57]. From a practical standpoint, these results suggest that analysis of MHC-I and -II expression levels could be developed to determine whether *CDK12* defective prostate tumors expressing higher levels of MHC genes are more likely to respond to ICB treatments.

Transcriptomic signatures associated with MHC expression variation have been described in different tumor types [37]. These signatures capture the activity of genes and biological pathways related to tumor cells’ crosstalk with the TME and appear to correlate with clinical responses to ICB [20,37,58–60]. Ayers *et al.* proposed a gene expression profile with eighteen genes relevant to predicting the clinical outcome of anti-PD1 therapy [58]. This IFN*-*γ gene signature in pretreatment tumor biopsies was associated with better results in melanoma, head-and-neck squamous cell carcinoma, and gastric cancer treated with pembrolizumab. In our study, the *CDK12* patient tumors expressing high levels of MHC genes showed the presence IFN*-*γ gene signature, which might reflect a better response to anti-PD1 therapy (Figure S9). Our results also showed the activation of pathways related to IFN*-*γ response in both the pPCA and mCRPC cohorts (Figure 1b, d, f, 3b, d, Figure 4b-c).

An IFN*-*γ gene signature can alternatively activate the expression of immunomodulatory mechanisms and promote adaptive resistance to therapeutic anti-PD1 [61,62]. IFN*-*γ is a central actor in the elimination phase during immunoediting [63]. However, exacerbated levels of IFN*-*γ are also associated with developing protumor molecular mechanisms leading to an immunosuppressive and tolerogenic TME [62]. We identified the upregulation of immunomodulatory molecules, such as *IDO1*, *TIM3,* and *CD274(PD-L1),* in both prostate tumors expressing higher levels of MHC-I and -II genes (Figure 2). Interestingly, the *CDK12* patient tumors expressing high levels of MHC genes showed the presence of IFN*-*γ gene signature, including upregulation of a non-classical MHC molecule (*HLA-E*), which is a known regulator of NK cell and CTL [64,65]. We also found a link between *IGHV* and *IGLV* over-expressed genes in a group of the *CDK12* defective pPCa expressing high levels of MHC-I (Figure 1a, upper cluster I). The production of Ig by tumor cells (cancer-derived Ig) is described in various tumors and PCa [66–69]. Also, cancer-derived Ig may act as checkpoint proteins and inhibit effector T cells and NK cells [67,70]. Our findings indicate the presence of an acquired resistance mechanism in a subset of *CDK12* defective prostate tumors expressing high levels of MHC genes. We suggest that measuring the basal expression of MHC genes could be developed as a biomarker to characterize *CDK12* patient samples with a previously inflamed environment and which would reflect the expression of immunomodulatory genes linked to resistance to anti-PD1.

The expression of MHC molecules plays a pivotal role in providing the signals necessary to recognize and activate the immune system against tumor neoantigens and is essential to controlling tumor growth through cytotoxic activity [63,71]. We found a higher abundance of CD8+ T cells in *CDK12* defective tumors expressing high levels of MHC genes (Figure 4). This result may either indicate the dependence of CD8+ T cells in MHC-I antigen presentation or suggest that enhanced CD4+ Th infiltrate could support the continued accumulation of CD8+ CTLs in the tumor microenvironment [59]. Interestingly, *CDK12* pPCa with high expression of both MHC-I and -II genes showed increased levels of γδ T cells (Figure 4a-b). The basal effector immune cell population, such as γδ T cells in pPCa, may contribute to IFN*-*γ signaling and indicate the dependence of the MHC-unrestricted recognition role of γδ T cells and its presence in the TME [20,72].

The loss of MHC expression could be derived from two different disruption means [73]. First, regulatory abnormalities downregulate the expression of MHC genes through mechanisms that do not affect the genomic structure of HLA genes. In such cases, specific T-cells can recover the MHC expression-mediated response (e.g., IFN-γ signaling). Second, during tumor evolution, the infiltration of cytotoxic lymphocytes eliminates tumor clones high immunogenic, causing a selection of clones that may present structural alteration, or “Hard” lesions, in the MHC loci or other genomic regions (e.g., *B2M*, *IFN*, *STAT*) distinct from the derived clone [48].

Genomic studies of chromosome 6 in various cancers suggest reduced MHC expression is often associated with genetic and LOH aberrations that may result in reduced antigen presentation and, thus, facilitate immune evasion [15,74]. In keeping with these data, our analysis of WES from pPCa and mCRPC revealed CN-LOH and LOH in *CDK12* defective tumors expressing low levels of MHC-I/II genes at known regulators of MHC expression and the components of the MHC (Table 2-3). Two pPCa patients also showed complete loss of the *B2M* locus, while one mCRPC exhibited a LOH event at the *B2M*, an important component of the MHC-I [48]. These results might indicate that after selection, the low expression of MHC genes is derived from tumor clones harboring “Hard” genomics alterations comprising loci involved in normal MHC expression. Therefore, these clones will be invisible to the immune system and contribute to tumor progression [18].

This study has some limitations. First, future studies in a large number of patients harboring *CDK12* mutations are needed to address potential statistical bias regarding our low number of patients. Second, we did not address the potential molecular mechanism underlying the alteration of the low expression of MHC in

*CDK12* defective PCa, including epigenetic alterations and miRNA activity. Further studies are needed to approach the causes of MHC disruption derived from “Soft” alterations that occur in *CDK12* defective prostate cancer. Third, because the non-tumor cell content was not defined in this study, we could not establish a limit for the contribution of the *CDK12* defective tumor cells and TME to the MHC expression since the RNA-seq relies on data from bulk tissue [75].

## Conclusion

We distinguished two sets of *CDK12* defective tumors based on differential MHC expression levels. Our data suggest that *CDK12* defective PCa expressing higher levels of classical MHC genes have an active and inflamed tumor microenvironment with elevated immunomodulatory pathway expression and increased presence of effector T cells. Tumors with decreased MHC expression showed copy-number alteration in genes that regulate MHC expression and antigen presentation. More extensive *in vitro* and *in vivo* investigations are required to relate this molecular subtype to potential actionable immunomodulatory mechanisms or possible therapeutic targets in PCa. Depending on the mechanism, the downregulation of MHC expression can sometimes be therapeutically restored to improve anti-tumor immunity [76]. These findings, therefore, support further investigations into the clinical use of MHC expression modulation in *CDK12* defective prostate cancer to identify tumors likely to respond to ICB treatment.

## List of abbreviations

*ADT*: Androgen Deprivation therapy
*APC*: Antigen-presenting cells
*CDK12*: Cyclin-dependent Kinase 12
*CN-LOH*: Copy-neutral loss of heterozygosity
*dbGaP*: The database of Genotypes and Phenotypes
*DDR*: DNA damage repair
*DEGs*: Differentially expressed genes
*FACETS*: Fraction and Allele-Specific Copy Number Estimated from Tumor Sequencing
*FDR*: False discovery rate
*GEP*: Gene expression profile
*GO*: Gene ontology
*ICB*: Immune-checkpoint blockade
*LM22*: Leukocyte gene signature matrix
*LOH*: loss of heterozygosity
*mCRPC*: Metastatic Castration-Resistant Prostate Cancer
*MHC*: Major histocompatibility complex
*MSigDB*: Molecular Signature Database
*NCCN*: National Comprehensive Cancer Network
*ORA*: Over-representation analyses
*PCa*: Prostate Cancer
*PE*: Paired-end
*pPCa*: Primary prostate cancer
*PRAD*: Prostate Adenocarcinoma
*PSA*: Prostate-specific antigen
*TCGA-PRAD*: The Cancer Genome Atlas
*TME*: Tumor microenvironment
*LOH*: Loss-of-Heterozygosity

## Ethics approval

This retrospective study was approved by the Ethics Committee in Research of Hospital of Ribeirão Preto, São Paulo, Brazil (HCRP) number CAAE 60032122.8.0000.5440 and the Ethics Board of the University of Toronto (Protocol: 00043323).

## Consent to publish

Not applicable.

## Availability of data and materials

The original dataset is deposited in the database of Genomic and Phenotypes (dbGaP) under accession numbers phs000178.v11.p8.c1, phs000915.v2.p2.c1. The original clinical information of both cohorts is deposited in the CbioPortal and GDC Portal. All other data supporting the conclusions of this article are included within the article and its additional files. Additional files related to our retrospective cohort are available at https://github.com/WilliamLautertD/CDK12_PCa.

## Authors’ contributions

Conception: W. L. D. and J. A. S

Interpretation or analysis of data: W. L. D., S. L. B., and J. A. S

Data acquisition, tissue handling, QC, performing assays: A.E.S., E.W., L.P.C., C.M.M., J.B. Preparation of the manuscript: W. L. D. and J. A. S.

Revision for important intellectual content: All authors Supervision: S. L. B. and J. A. S.

## Acknowledgments

The authors acknowledge the contributions of the Department of Surgery and Anatomy, the Department of Pathology, the Medicine School of Ribeirao Preto, and members of Diagnostic Development at the Ontario Institute for Cancer Research. The authors are grateful to L. A. F. B. and S. P. F. B. for the informatic technical support in the early stage of the project, Genomic and Molecular laboratory/PUCRS for technical assistance, and dbGap and the Cbioportal for archiving and distributing the data from the above-cited studies.

