## Additional file 1 for "Quantitative MHC class-I/-II gene expression levels in *CDK12* mutated prostate cancer reveal intratumorally T cell adaptive immune response in tumors"

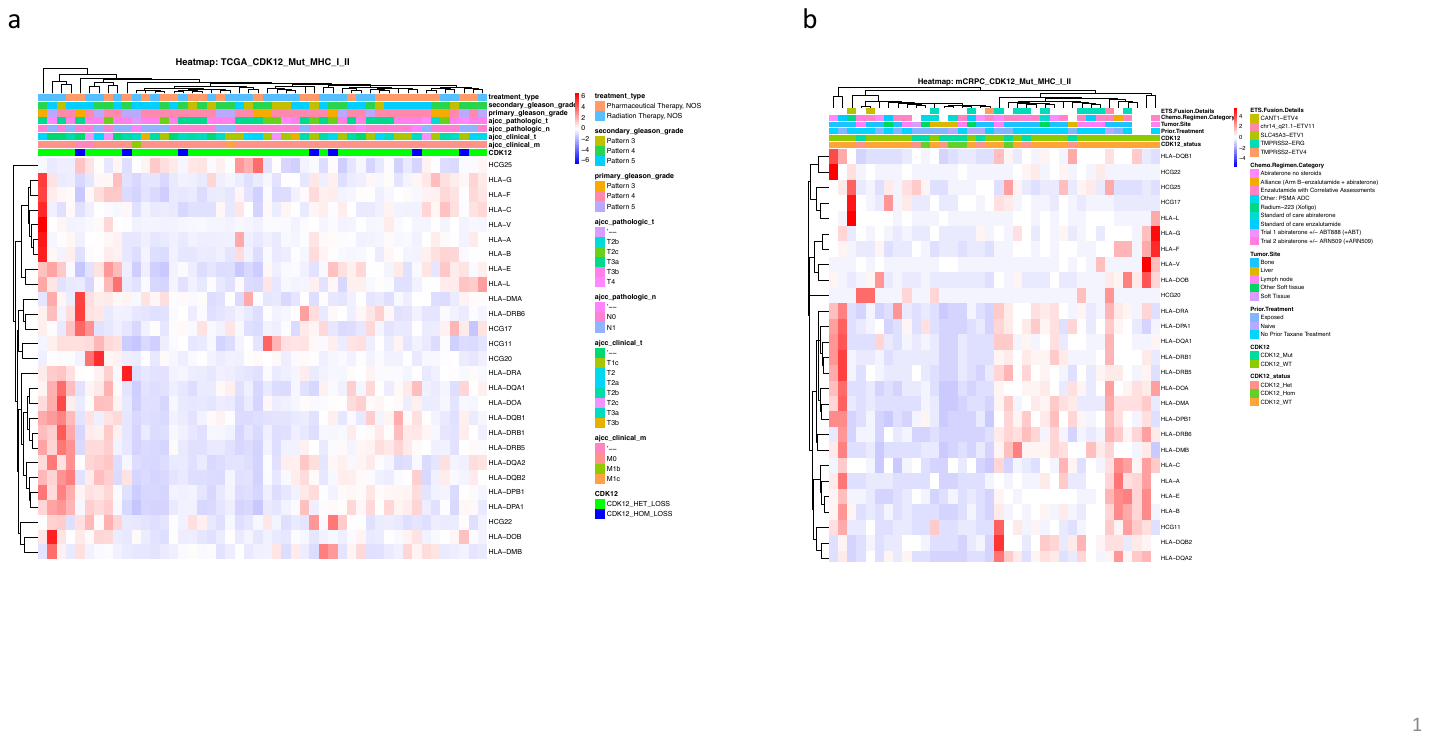

**Figure S1. HLA expression in *CDK12* patients.** We observed two groups of *CDK12* defective tumors regarding MHC expression using hierarchical clustering analysis for pPCa (a) and mCPRC (b). We then used the normalized expression level of each gene to classify the patients based on their MHC gene expression (See method and Fig. S2). The summary values used to classify the tumors are shown in table S1.

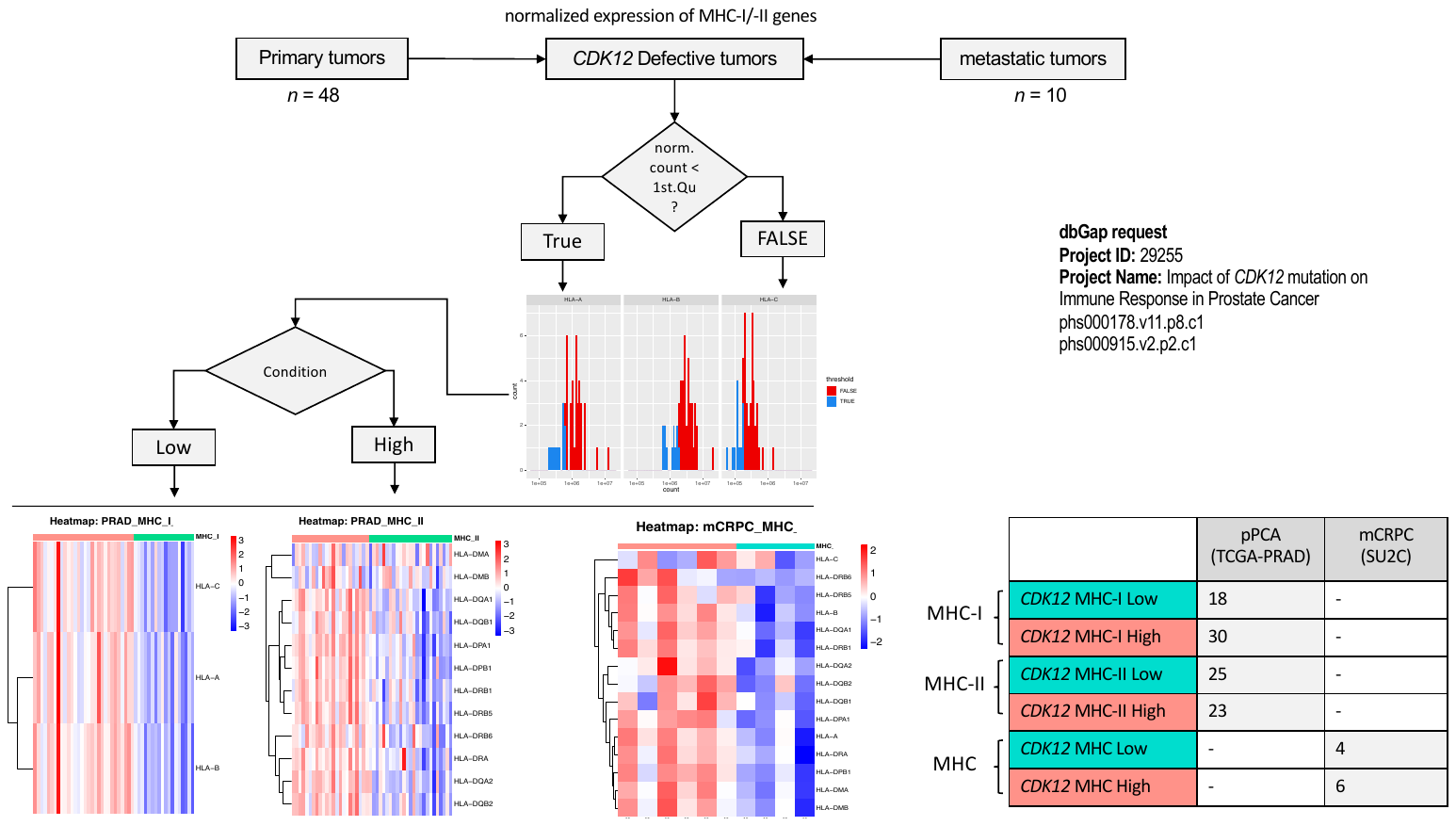

**Figure S2. Classification of *CDK12* patients based on HLA genes (MHC-I and -II) expression.** We used the normalized expression values of the classical genes that composed each MHC class (e.g., MHC-I: *HLA-A*, *HLA-B*, *HLA-C*; MHC-II: *HLA-DPA1*, *HLA-DPB1*, *HLA-DQA1*, *HLA-DQB1*, *HLA-DQB2*, *HLA-DRA*, *HLA-DRB5*, *HLA-DRB6*) (Schaafsma et al. 2021). We dichotomized expression levels below or above the first quartile for each gene using the normalized read count; we then used this classification to generate the final logical values regarding the patient’s MHC status (‘High’ or ‘Low’ expressed) as follows: e.g., threshold *Gene 1* **AND** threshold *Gene 2* **AND** threshold *Gene 3*. Samples were classified as having MHC ‘Low’ expression when at least one gene composing each class was expressed at a low level. In mCRPC *CDK12*- defective, all tumors classified as MHC-High presented similar high expression of both MHC-I and -II; thus, we called these tumors just MHC High. The table above shows the final number of patients classified for each tumor type in each condition. The summary values used to classify the tumors are shown in table S1. The final logical values are shown in additional tables 3, 4, 5, and 6.

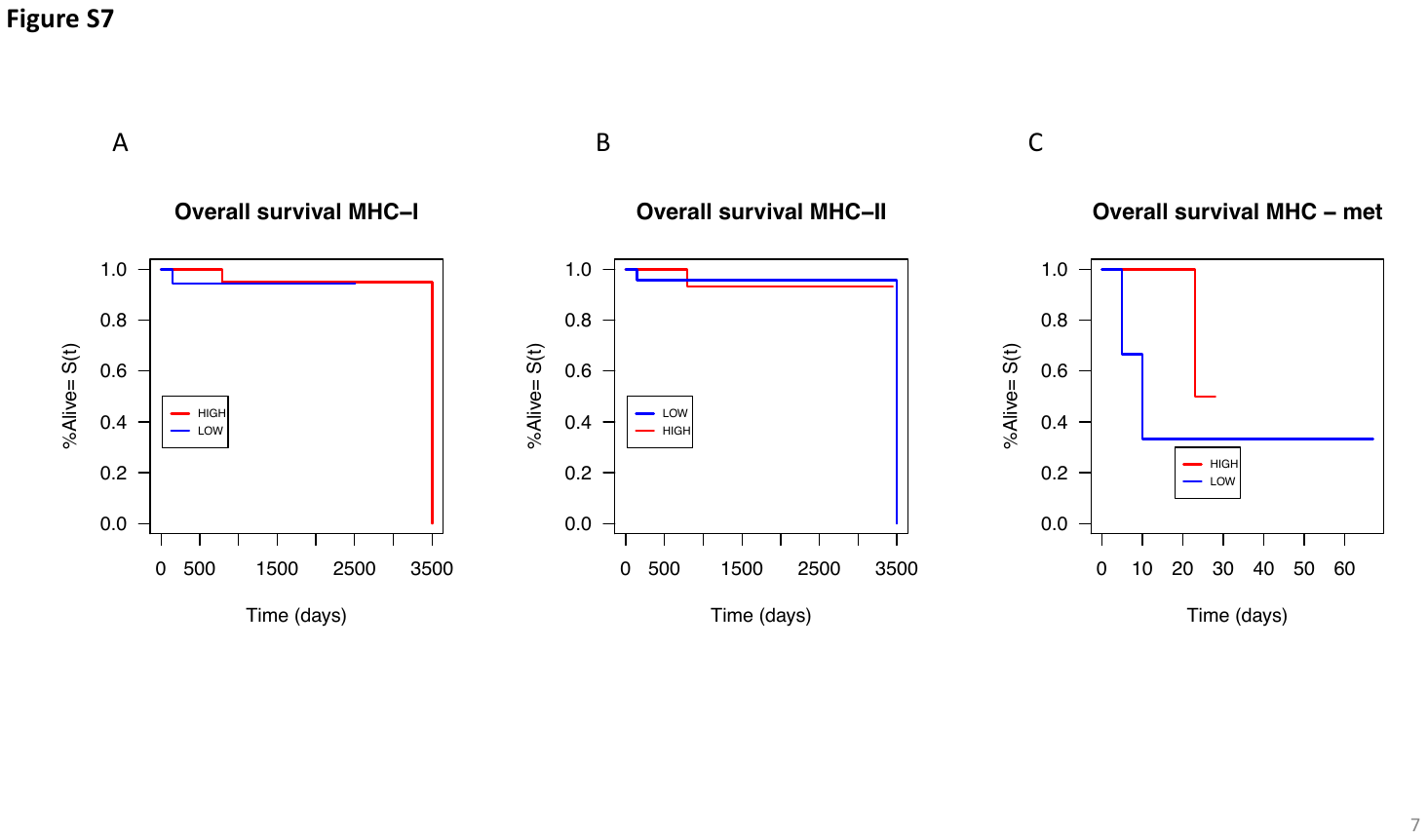

**Figure S3. Overall survival of CDK12 defective patients. (A)** Kaplan-Maier curve for overall survival in CDK12 defective MHC-I High expressed primary tumors, **(B)** Kaplan-Maier curve for overall survival in CDK12 defective MHC-II High expressed primary tumors, and **(C)** Kaplan-Maier curve for overall survival in CDK12 defective MHC High expressed metastatic tumors. Kaplan-Maier curve for overall survival was calculated using the *survfit* function from the survival package for R. The difference was not significant in all the comparisons (*p* = 0.6, *p* = 1, and *p*= 0.5, respectively). Log-rank statistic tests were performed using the *survdiff* function.

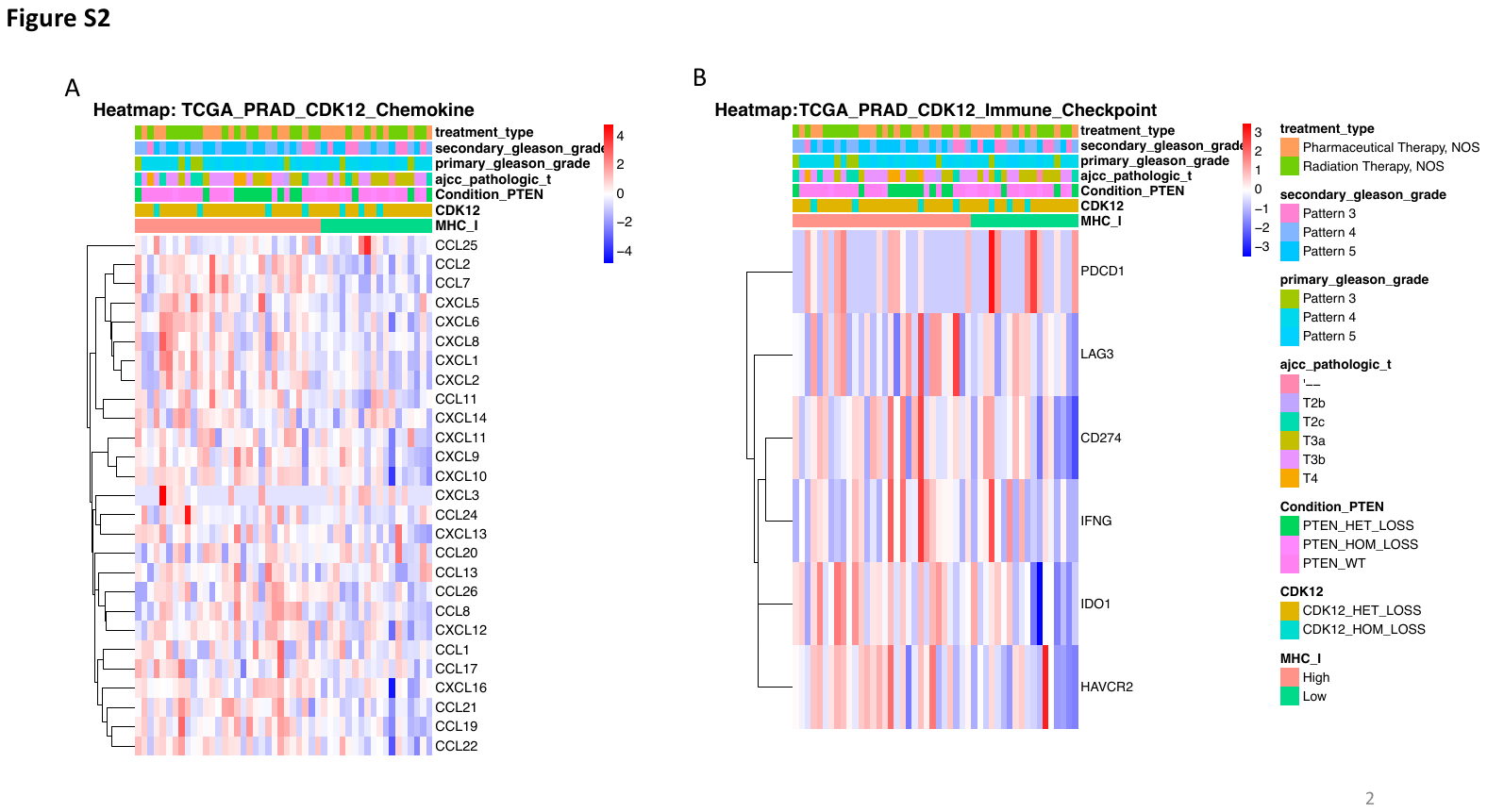

**Figure S4. Expression profile of the tumor microenvironment of primary prostate tumors**. Supervised heatmap of chemokine expression (a) and supervised heatmap of immunomodulatory gene expression in the pPCa *CDK12*-Mut expressing high levels of *MHC-I* (b). Clinical information is displayed on top of each patient. Color scale represents the Z-score of the normalized read counts for each gene, where red scale color approaching red means high expressed and blue low expressed genes. n = 48 (CDK12-Mut MHC-I High = 30, CDK12-Mut MHC-I Low = 18).

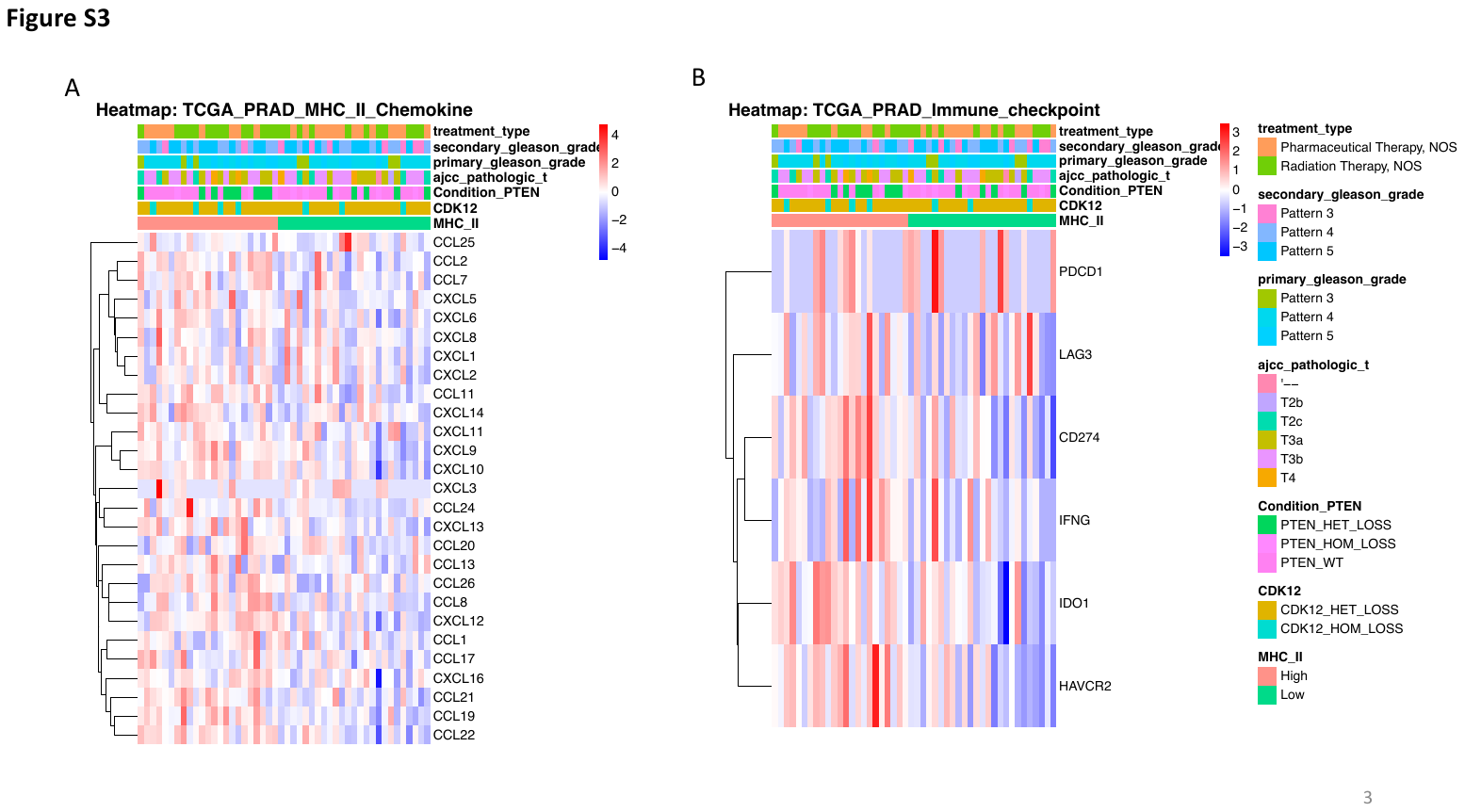

**Figure S5. Expression profile of the tumor microenvironment of primary prostate tumors.** Supervised heatmap of chemokine expression (a) and supervised heatmap of immunomodulatory gene expression in the pPCa *CDK12*-Mut expressing high levels of *MHC-II* (b). Clinical information is displayed on top of each patient. Color scale represents the Z-score of the normalized read counts for each gene, where red scale color approaching red means high expressed and blue low expressed genes. n = 48 (CDK12-Mut MHC-II High = 23, CDK12-Mut MHC-II Low = 25).

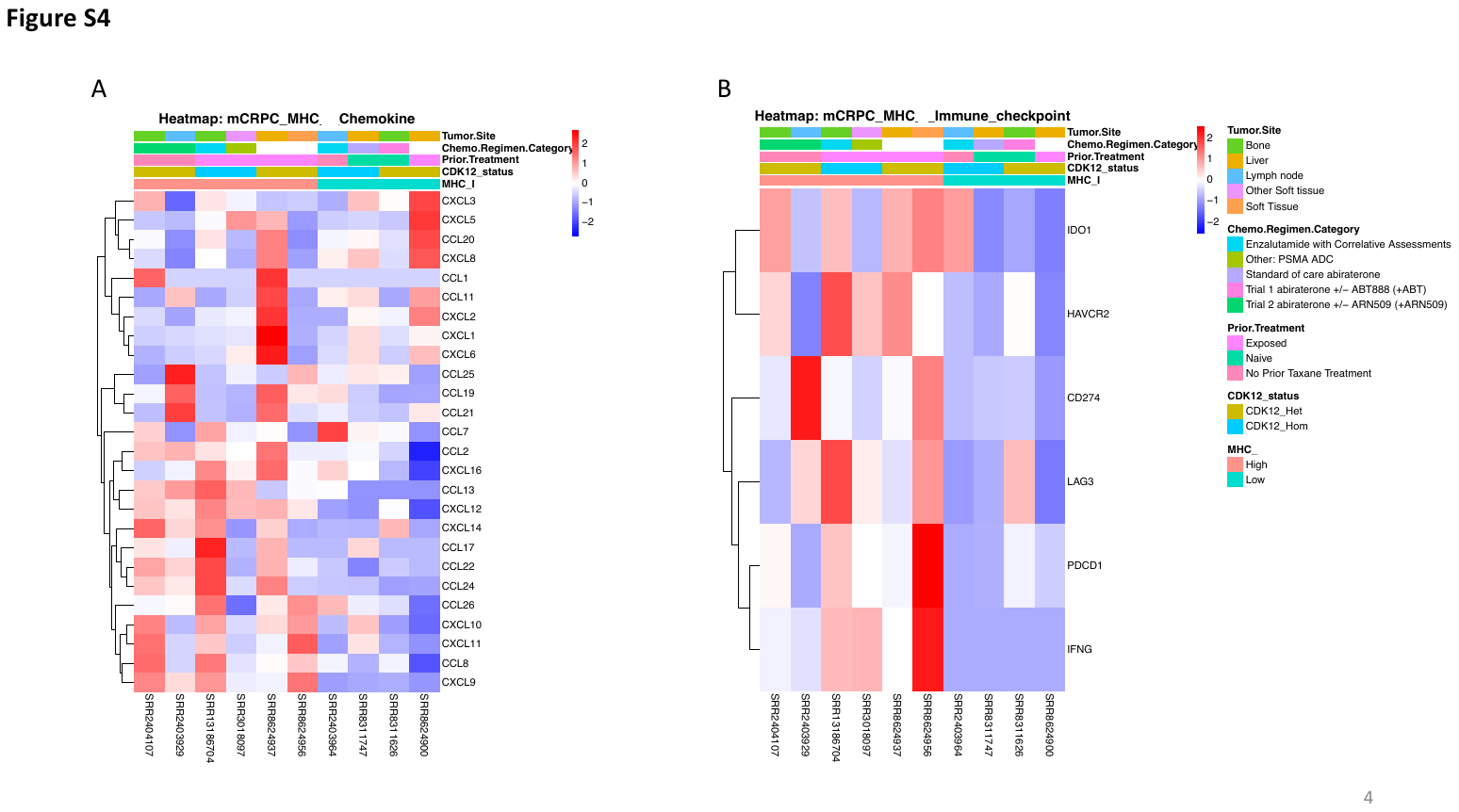

**Figure S6. Expression profile of the tumor microenvironment of metastatic castration-resistant prostate tumors.** Supervised heatmap of chemokine expression (a) and supervised heatmap of immunomodulatory gene expression in the mCRPC *CDK12*-Mut expressing high levels of *MHC* genes (b). Clinical information is displayed on top of each patient. Color scale represents the Z-score of the normalized read counts for each gene, where red scale color approaching red means high expressed and blue low expressed genes. n = 10 (CDK12-Mut MHC High = 6, CDK12-Mut MHC Low = 4).

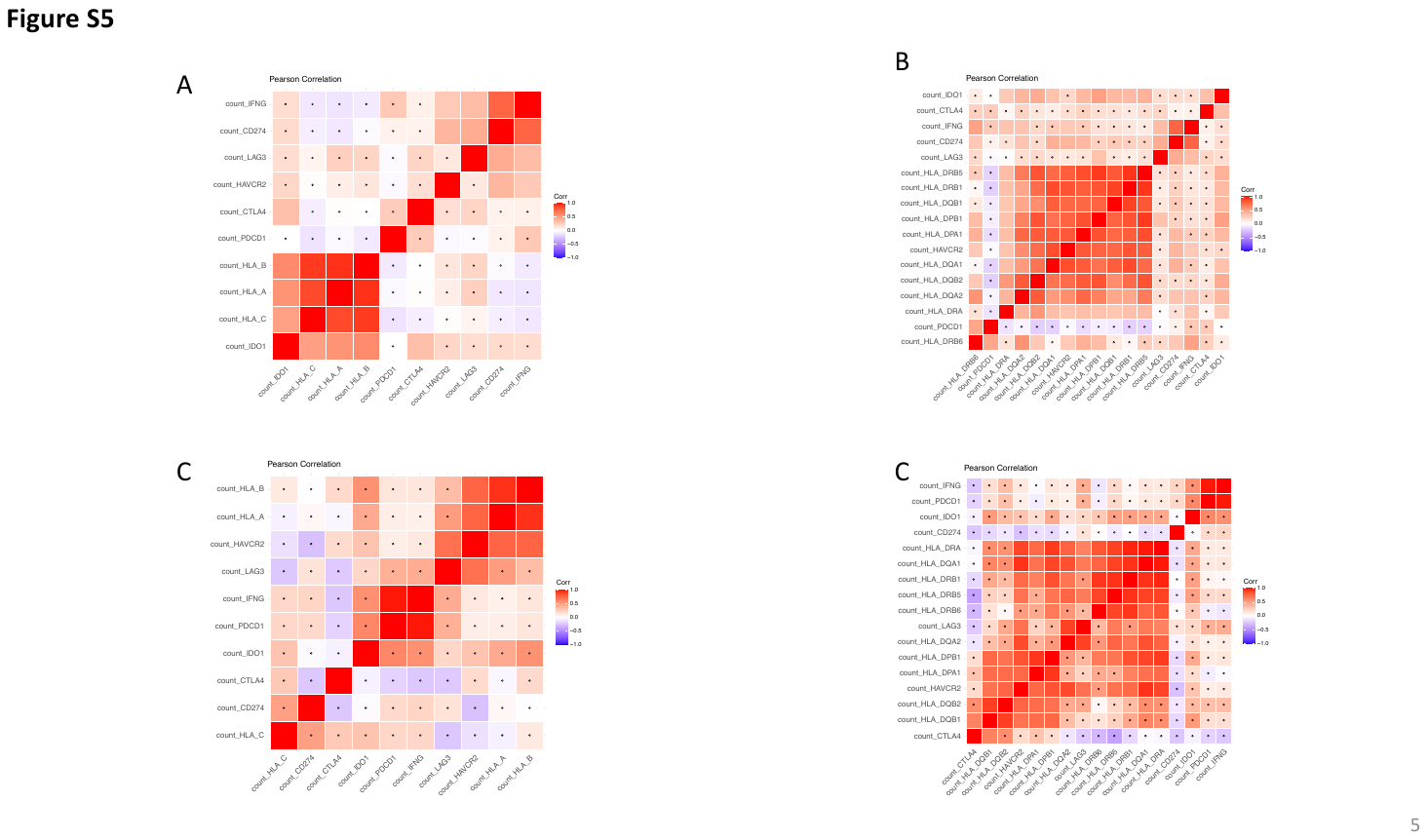

**Figure S7. Correlation of classical MHC and immunomodulatory genes.** (a-b) Correlation plots between *MHC-I /–II* and the immunomodulatory genes in primary prostate tumors. (c-d) Correlation plots between *MHC-I /–II* and the immunomodulatory genes in metastatic castration-resistant prostate tumors. Pearson Correlation was used in normalized expression levels of each gene (coef. level = 0.95). A significant correlation is indicated in nonmarket squares (*p*<0.05).

**Figure S8.** **Whole-genome sequencing data from *CDK12* defective in primary MHC low-expressed tumors.** Integrated visualization shows the total copy number log ratio (logR) on the top panel, and allele-specific log-odds-ratio data (logOR) on the second panel with chromosomes alternating in blue and gray. The third panel plots the corresponding integer (total, minor) copy number calls. Tumors with a 2:1 ratio are considered normal for each position, while 2:0 and 1:0 represent CN-LOH and LOH events. The estimated cellular fraction profile is plotted at the bottom, revealing both clonal and subclonal copy number events.

**Figure S9.** **Whole-genome sequencing data from *CDK12* defective in metastatic MHC low-expressed tumors.** Integrated visualization shows the total copy number log ratio (logR) on the top panel, and allele-specific log-odds-ratio data (logOR) on the second panel with chromosomes alternating in blue and gray. The third panel plots the corresponding integer (total, minor) copy number calls. Tumors with a 2:1 ratio are considered normal for each position, while 2:0 and 1:0 represent CN-LOH and LOH events. The estimated cellular fraction profile is plotted at the bottom, revealing both clonal and subclonal copy number events.

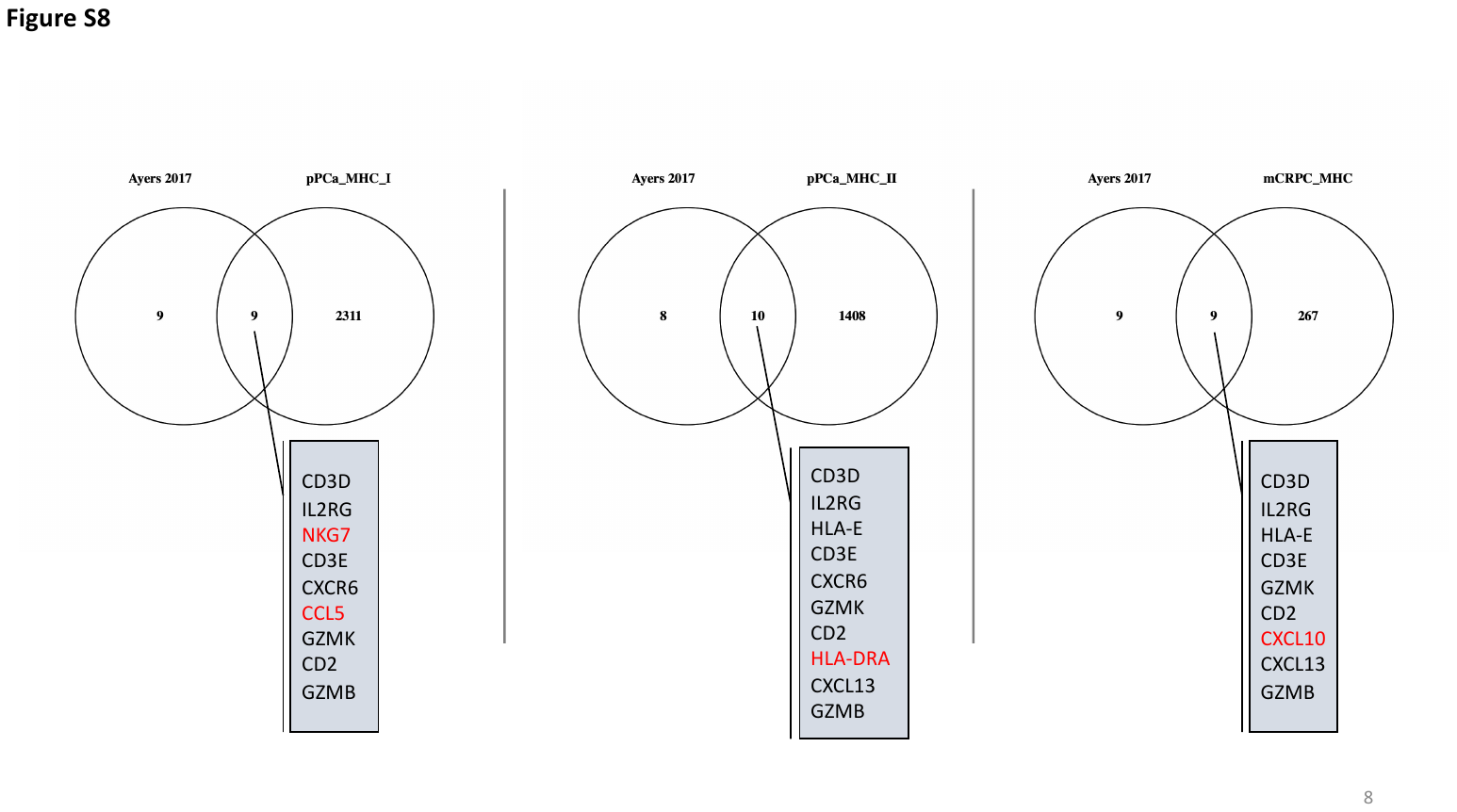

**Figure S10.** **Common activation of IFN-γ-responsive genes in *CDK12* defective MHC high expressed tumors.** In our study, the *CDK12* patient tumors classified as MHC High showed the presence of several genes previously described as IFN-γ-responsive and cytotoxic activity and associated with response to anti-PD1 therapy (Ayers, 2017. <https://doi.org/10.1172/JCI91190>). Common genes with Ayers et al. 2017, are indicated in the intersection and unique features for each comparison are indicated in red. Venn`s Diagrams by *Oliveros, J.C. (2007-2015) Venny. An interactive tool for comparing lists with Venn's diagrams.* <https://bioinfogp.cnb.csic.es/tools/venny/index.html>.
